# Conspicuous shell shape plasticity across lake-stream habitats in a freshwater mussel (*Pyganodon grandis*)

**DOI:** 10.1101/2023.10.30.564758

**Authors:** Sean M. Keogh, Ben J. Minerich, Lindsay M. Ohlman, Madeline E. Pletta, Anna E. Scheunemann, Zoe K. Schroeder, Zebulin A. Secrist, Alex J. Franzen, Bernard E. Sietman, Andrew M. Simons

## Abstract

2.

Hydrodynamic forces and their absence appears to exert differential selection pressure between lake and stream populations, creating a tight fit between organismal phenotypes and their environments. Ecophenotypic variants may be the result from isolated genetic evolution or phenotypic plasticity, where a widespread genotype can produce multiple phenotypes dependent on the environment. Freshwater mussels possess a wide degree of morphological variation that frequently covaries with the environment, making them a good system to understand the mechanisms of ecophenotypic variation across hydrological conditions. We designed a two-year experiment where individuals from the same *Pyganodon grandis* maternal brood (half-full siblings) were reared at a controlled site and four natural sites involving one lake and three streams. At the end of the experiment, shell shape was quantified for recaptured (N=70), wild (N=206), and zoo-reared (N=305) mussels. Analysis of covariance found significant differences in shell shape between rearing sites, particularly between stream and lake habitats, but no shape differences were detected across the three stream sites. At two of the four sites, shell shape of recaptured individuals matched the morphology of wild populations. Genomic sequencing and parentage analysis identified 27 different fathers among recaptured individuals. Yet, no genetic differences were present between stream and lake habitats and there was no effect of parentage on shell shape. Taken together, phenotypic plasticity, over genetic differentiation, is identified as the primary mechanism of shell shape ecophenotypy. Plasticity may be a key adaptation for freshwater mussels and possibly provide a buffer against their imperilment in degraded habitats.

**Summary statement:** Streamlined and obese freshwater mussel shell shapes in stream-lake environments are developed from the same maternal brood and may reflect adaptations to the presence and absence of streamflow.

## 3. Introduction

Understanding the role in which extrinsic, abiotic forces shape phenotypic biodiversity is fundamental to understanding biological diversification. When species exist across heterogeneous environments, evolutionary processes can produce localized phenotypic variants creating a tight phenotype-environment fit within species, which may reflect adaptation to local conditions (Hendry 2016; Kawecki and Ebert 2004). Two major pathways exist to explain this pattern: 1) localized genetic evolution across populations in different habitats can result in genotypic differentiation (Kawecki and Ebert 2004), producing divergent phenotypes and 2) ecophenotypic variation can be the result of phenotypic plasticity, where a single genotype produces different phenotypes dependent on environmental cues (Pigliucci 2005; West-Eberhard 2003).

Aquatic organisms that exist across hydrological gradients, like between lake and stream environments, must cope with the presence or complete absence of streamflow. This continuous hydrodynamic force, or lack thereof, appears to exert differential selection pressure between lake and stream populations, creating body shape and gill raker divergences in fishes (Oke et al. 2016; Berner et al. 2008; Samways et al. 2015; Theis et al. 2014; Cureton and Broughton 2014), gill morphological differences in conspecific caddisflies (Parisek et al. 2023), and leaf shape divergence in aquatic plants (Ganie et al. 2015). Identifying the mechanism of such lake-stream phenotypic divergences requires experimentation, typically through common garden (organisms from different environments reared in the same environment) or reciprocal transplant (organisms reared in one environment then transferred to another environment). By controlling the environment, these approaches measure phenotypic differences or the ‘reaction norm’ either between individuals (common garden) or between environments (reciprocal transplant) thus separating the environmental versus genetic effects on phenotypic variation (Hendry 2016; Pfennig 2021; Whelan 2021). A similar but more strict approach standardizes genetic variation through genetic clones or siblings. The reaction norms of clones or family groups can then be measured across environmental variation and then compared to wild populations in shared environments to not only separate genetic versus plastic effects but assess their relative roles in ecophenotypic outcomes.

Like other aquatic lineages, many freshwater mussels (Family Unionidae) possess intraspecific phenotypic variation across lake and stream environments (e.g. *Amblema plicata*, *Eurynia dilatata*, *Fusconaia flava*, *Lampsilis siliquoidea*, *Ligumia recta*, *Pleurobema sintoxia*, and *Pyganodon grandis*) (Ortmann 1919; Grier 1920). These ecophenotypic patterns are expressed across divergent lineages (Ortmann 1920) and therefore are assumed to be adaptive, specifically in habitat specific functions: shell compression (decreased shell width) for stability against streamflow and shell inflation (increased shell width) for buoyancy or metabolic optimization in lake environments (Haag 2012; Watters 1994). Importantly, an underlying process explaining this ecophenotypic pattern is unknown. Many integrative studies using morphometric, molecular phylogenetic, and/or population genetic tools have failed to find genetic correlation with ecophenotypic variation (Baker et al. 2003; Cyr et al. 2007; Inoue et al. 2013; Olivera-Hyde et al. 2020; Zieritz et al. 2010) leading to phenotypic plasticity as the default explanation for all intraspecific morphological variation in freshwater mussels (Inoue et al. 2013; Zieritz et al. 2010; Hornbach et al. 2010; Jeratthitikul et al. 2019; Sheldon 2017; Wu et al. 2022). Although these molecular studies provide compelling evidence supporting phenotypic plasticity, they do not sequence markers involved in shell production and thus genetic mechanisms (e.g. convergent evolution) cannot be ruled out. An experimental approach that standardizes the environment, genetic variation, or both is needed to separate genetic differentiation from phenotypic plasticity. Freshwater mussels are among the most imperiled aquatic groups globally and understanding if ecophenotypes are the result of speciation, genetic differentiation, or phenotypic plasticity is critical to successful conservation management (Ferreira-Rodríguez et al. 2019).

Here, we conducted a field experiment to understand the mechanism(s) of ecophenotypic variation in freshwater mussels. We performed the experiment using *Pyganodon grandis*, a species with a high growth rate relative to other freshwater mussels (Kesler and Van Tol 2000) and is easily propagated in the laboratory (M. Bradley, pers. comm.). *Pyganodon grandis* shares the ecophenotypic pattern of shell inflation in lakes and compression in streams but additionally becomes more circular in shell outline in lakes and more elongated in streams (Haag 2012) (Figure 1a). Ecophenotypic variation is so dramatic, conspecifics were historically recognized as separate species or subspecies (Baker 1928). We used experimental mussels reared from the same maternal brood, to minimize genetic variation, in one of two sets of wild common gardens: stream and lake habitat types comprising multiple field sites per habitat type. We asked six related questions: among recaptured experimental mussels, (1) does morphology differ between rearing sites, (2) habitat type, or (3) between sites with the same habitat type? Additionally, to assess the potential bounds of plasticity we compared experimental mussels to wild populations to ask (4) to what extent does recaptured mussel morphology match the morphology of wild *P. grandis* reared within the same site and (5) habitat type? Lastly, we used genomic markers to ask (6) does paternal parentage influence phenotypic outcomes?

**Figure 1:**
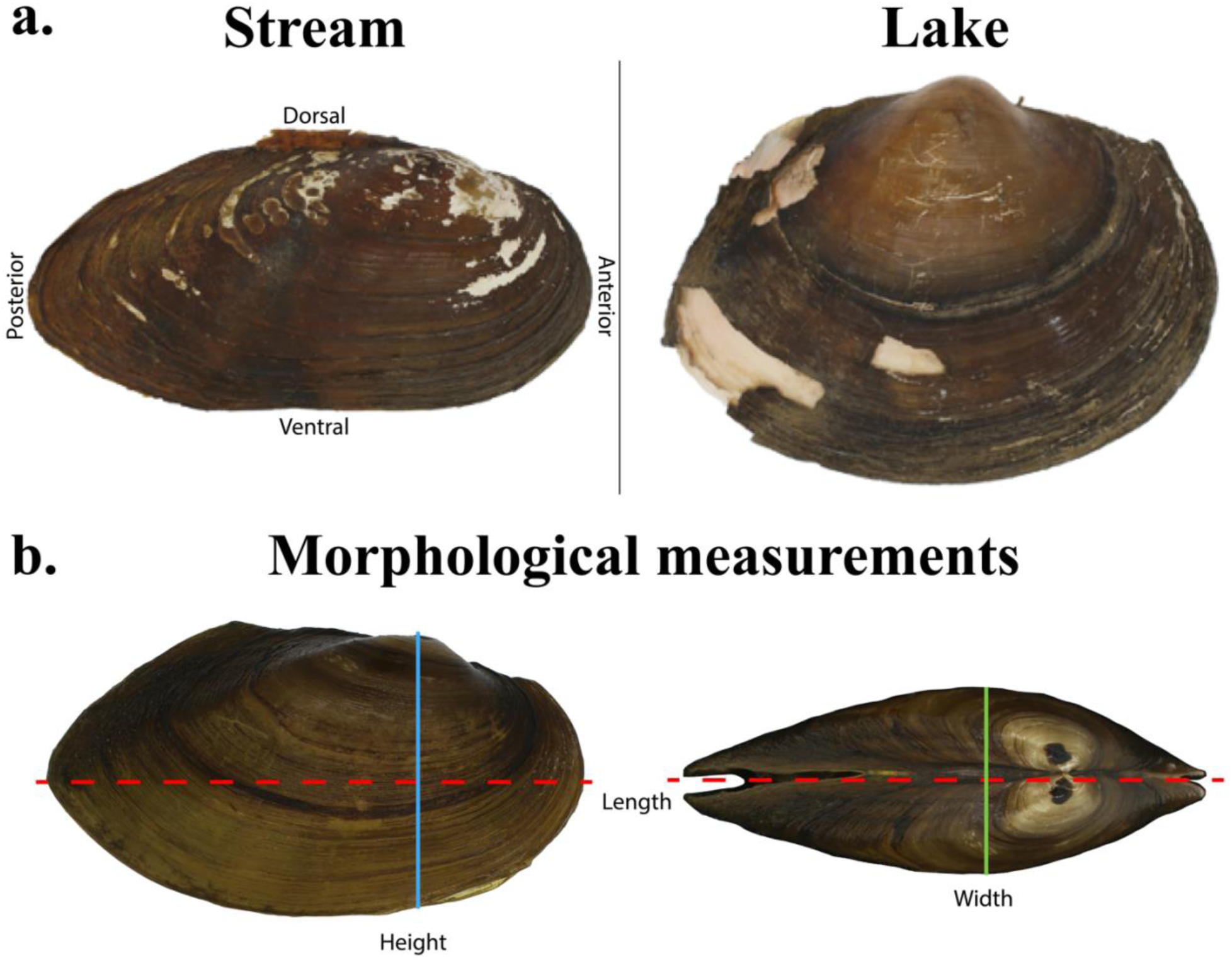
(a.) Typical shell shape variation of *P. grandis* between stream and lake habitats (Ortmann 1919; Grier 1920; Haag 2012). (b.) Morphological measurements shell length, height, and width (mm) quantified in this study.

## 4. Materials and Methods

### 4.1 Experimental subjects and design

We collected a gravid female *P. grandis* from Chub Creek in southeastern Minnesota, USA (Cannon River drainage; 44.522, –93.2) on April 25, 2020. The gravid female was promptly transported to the Minnesota Department of Natural Resources Center for Aquatic Mollusk Programs and held in an aerated container for four days. *Pyganodon grandis*, like most freshwater mussels, is an obligate parasite on freshwater fishes during the larval (hereafter ‘glochidial’) life stage and can parasitize multiple host species (Clarke and Berg 1959; Penn 1939; Trdan and Hoeh 1982; Wilson 1916). However, to eliminate possible host effects on morphological variation of *P. grandis*, we used a single host, *Perca flavescens* (Yellow Perch), to rear mussels. Eighteen *P. flavescens* were manually inoculated with glochidia from the collected *P. grandis* individual using standard freshwater mussel-fish host inoculation protocols (Zale and Neves 1982; Sietman et al. 2018). Briefly, viable glochidia were manually extracted and suspended in aerated water baths with *P. flavescens* hosts. We visually inspected the gills and fin margins of hosts to confirm glochidium attachment and subsequently moved fish to recirculating aquaria fitted with a collection net at each outflow. We monitored contents of nets regularly for sloughed glochidia and transformed juvenile mussels. Juvenile mussels began dropping off seventeen surviving fish twelve days after inoculation and continued for ten days (May 11-20, 2020). We aggregated ∼14,765 juveniles in aerated buckets and transferred metamorphosed mussels to the Minnesota Zoo mussel propagation facility. Mussels were held in mesh-lined (400um) baskets with sand (grain size 0.35-0.45mm) in a flow-through trough and fed a commercial algae diet (Nanno 3600 and Shellfish Diet 1800) along with filtered (90um screen) lake water.

Three months later (August 2020) mussels were 8-17mm in length and could be safely marked. We marked both valves of approximately 5,860 mussels with a dot of black cyanoacrylate glue cured with an accelerant (StewMac Inc. Athens, OH, USA). We found nine suitable release sites (four streams, five lakes) in the Cannon River drainage (aside from one lake site just south of the confluence of the Cannon and Mississippi Rivers) and in close proximity to the collection site of the mother *P. grandis* (Figure 2). We randomly selected batches of 420-1,500 mussels to release at each of the nine sites over a five-week period (mid-August-September). In addition, ∼900 mussels were split between two mesh-lined baskets suspended in a lake behind the zoo’s propagation facility (same water source used to feed the reared mussels). Mussels were transported to release sites in five-gallon buckets and acclimated to release sites with a 50% water change over 30 minutes. At deeper lake sites, we set up ten meters of lead line spread between PVC pipes and a cinder block between them to landmark release locations. At stream sites, we scattered mussels over approximately 100 meters of stream distance.

**Figure 2:**
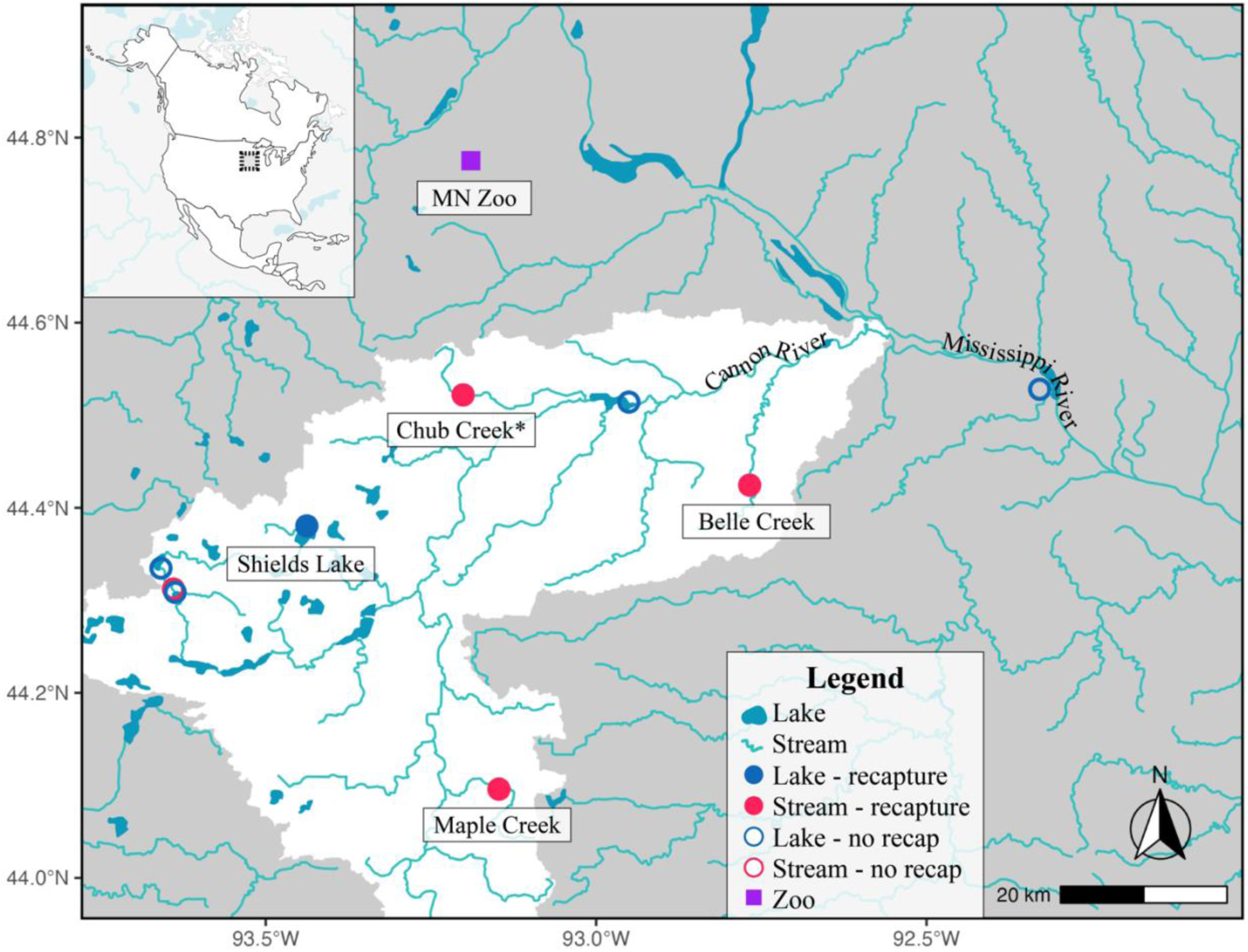
Map of the Cannon River drainage in southeastern Minnesota, USA (white). Circular points denote release sites: hollow = no recapture, solid = recaptures. *Denotes the location of the gravid female’s field collection (Chub Creek).

### 4.2 Recapture and data collection

Two years following release (August 2022), we returned to field sites and attempted to recapture released individuals using standard field collecting methods (e.g. wade, snorkel, SCUBA) over a three-week period. Recaptured individuals were immediately placed on ice and transported to the University of Minnesota where they were phenotyped, a tissue sample was taken, and then stored in a ultracold freezer (–80C) with the rest of the animal. We phenotyped each individual using three measurements frequently used in freshwater mussel morphometrics with a digital caliper (Mitutoyo Canada Inc. Mississauga, ON, CA): maximum length (mm), maximum height (measured perpendicular to length centered on umbo; mm), and maximum width (mm) (Figure 1b). Individuals that were kept at the Minnesota Zoo were measured similarly. At all sites we successfully recaptured (marked) mussels, we also collected and quantified the morphology of wild populations of *P. grandis*. Collection of wild individuals (primarily live individuals but some empty shells) occurred both at time of release and recapture. However, only vouchered specimens were measured during time of release to prevent pseudo replication.

### 4.3 Genomic sequencing, bioinformatics, and paternity testing

Although experimental subjects were reared from the same maternal brood and thus were at least half siblings, multiple paternity has been discovered in many freshwater mussel species (Bai et al. 2012; Christian et al. 2007; Ferguson et al. 2013; Garrison et al. 2021; Inoue et al. 2023; Wacker et al. 2018; 2019) and may play a role in phenotypic expression. Therefore, we generated single nucleotide polymorphisms (SNPs) from 3RAD-sequencing (Bayona-Vásquez et al. 2019) of 46 recaptured mussels and the maternal mussel to identify paternal parentage and its effect on progeny shell shape. Briefly, we extracted genomic DNA from the foot and/or mantle tissue using the Qiagen DNAeasy Blood and Tissue Kit (Valencia, CA, USA) and quantified DNA using a Qubit Fluorometer (Thermo Fisher Scientific; Waltham, MA, USA). DNA was shipped to the EHS DNA Laboratory at University of Georgia. Samples were normalized then digested with enzymes XbaI, EcoRI, and NheI followed by iTru adapter ligation of variable length internal indexes. 3RAD libraries were amplified by PCR with iTru5 and iTru7 primers (Glenn et al. 2019). A Pippin Prep (Sage Science, Inc. Beverly, MA, USA) was used to size select DNA fragments and sequenced at Novogene Inc. (Sacramento, CA, USA) on an Illumina HiSeq X Ten (San Diego, CA, USA). Raw 3RAD reads were inspected for quality in FastQC v0.11.7 (Andrews 2010). Seven samples had low quality reads and were removed from the assembly. We assembled reads in ipyrad v0.9.92 (Eaton and Overcast 2020) with default settings aside from the following modifications: assembly method set to “denovo”, datatype “pair3rad”, restriction overhang “GCTAG, TAATTC”, clustering threshold “0.88”, and minimum number of samples per locus (missing data parameter) “20” (∼50% missing). This resulted in a cumulative 33,628 SNPs across 39 samples.

We used the VCF output file from ipyrad and VCFtools v0.1.16 (Danecek et al. 2011) to extract 355 SNPs that were unlinked, biallelic, had a call rate >90%, and minor allele frequency >0.03. We used the R package *dartR* (Gruber et al. 2018) to calculate genetic distances (F_ST_) between rearing sites and habitats. Finally, we used COLONY (Jones and Wang 2010) to estimate paternal sibship. The VCF file was converted to a COLONY input format in the R package *radiator* (Gosselin et al. 2020). Long, full likelihood runs were performed with five replications. Because all sequenced individuals (aside from the mother) were from the same maternal brood we selected polygamous mating system for females, monogamous mating system for males, and all offspring as (at least) half siblings with a known mother.

### 4.4 Statistical analysis

All statistical analyses were performed in R Statistical Environment v4.3.1 (R Core Team 2023) using packages *tidyverse* (Wickham et al. 2019), *car* (Fox and Weisberg 2019), and *multcomp* (Hothorn et al. 2008). We ran linear regressions predicting shell height or width dependent on length. Regressions were run for each site and habitat type. Additionally, we constructed ANCOVA (analysis of covariance) models to test a series of questions below. We used type II sums of squares and α ≤ 0.05 was used to assess significance. For each model, shell height or width was used as a dependent variable with shell length as a covariate and the interaction of shell length and the grouping variable of interest.

#### 4.4.1 Question 1: Do mussels reared at different sites have different shell morphology?

A first order analysis to test if the environment has an effect on shell morphology was to perform ANCOVAs to test if rearing sites had significantly different shell morphology (size relative height and width). Only recaptured mussels were included but models were constructed with and without individuals kept at the zoo.

#### 4.4.2 Question 2: Do mussels reared at different habitats have different shell morphology?

We wished to test what environmental factor(s) were causing any morphological differences between rearing sites. Therefore, we aggregated recaptured mussels reared at different sites but in the same habitat, stream and lake, and tested for a significant effect of habitat type on shell morphology. As with question 1, only recaptured mussels were included. Although the zoo reared individuals were technically reared in a lake environment, they were in a densely packed enclosure with flowing, aerated water, and therefore were excluded from subsequent analyses.

#### 4.4.3 Question 3: Do mussels reared at different sites but the same habitat type have different shell morphology?

To test if rearing site had an effect on shell morphology among mussels reared in comparable habitats, we constructed ANCOVA models consisting of only stream or lake reared individuals with site used as the predictor variable. We then used pairwise *post hoc* testing (Tukey method) to assess significant differences between each site comparison. Because only one site was a lake environment, this test was only done for stream sites.

#### 4.4.4 Question 4: Do recaptured mussels have different shell morphology than wild mussels reared at the same site?

If shell shape is purely determined by environmental factors in *P. grandis*, then recaptured mussel morphology should match the morphology of wild individuals reared at the same site. Therefore, we aggregated recaptured and wild collected mussels at each site and tested for morphological differences.

#### 4.4.5 Question 5: Do recaptured mussels have different shell morphology than wild mussels reared at the same habitat type?

We combined all stream reared mussels across the three stream sites and tested for morphological differences between recaptured and wild mussels. Because only one site was a lake environment, this test was only performed for stream sites.

#### 4.4.6 Question 6: Does paternity affect shell morphology?

The COLONY analysis allowed us to assign each sequenced individual to an unsampled father. We then tested for paternal effects on shell shape while using habitat type as a covariate in ANCOVA models because of its strong effect on shell shape.

## 5. Results

### 5.1 Recaptured and wild caught mussels

We recaptured mussels from three of four stream sites and only one of the five lakes (Figure 2-3). Recapture sites included Belle Creek (44.424, –92.768), Chub Creek (44.522, – 93.2), Maple Creek (44.0958, –93.15), and Shields Lake (44.37, –93.43). The Chub Creek recapture site was also where the mother was collected. In total, 70 mussels were recaptured including 26 from Shields Lake and 44 from the three stream sites. These samples exclude the 305 experimental control mussels kept at the zoo for the duration of the experiment. In addition to recaptured mussels, a cumulative 206 wild *P. grandis* were collected from the same sites (Supplementary Table 1).

**Figure 3:**
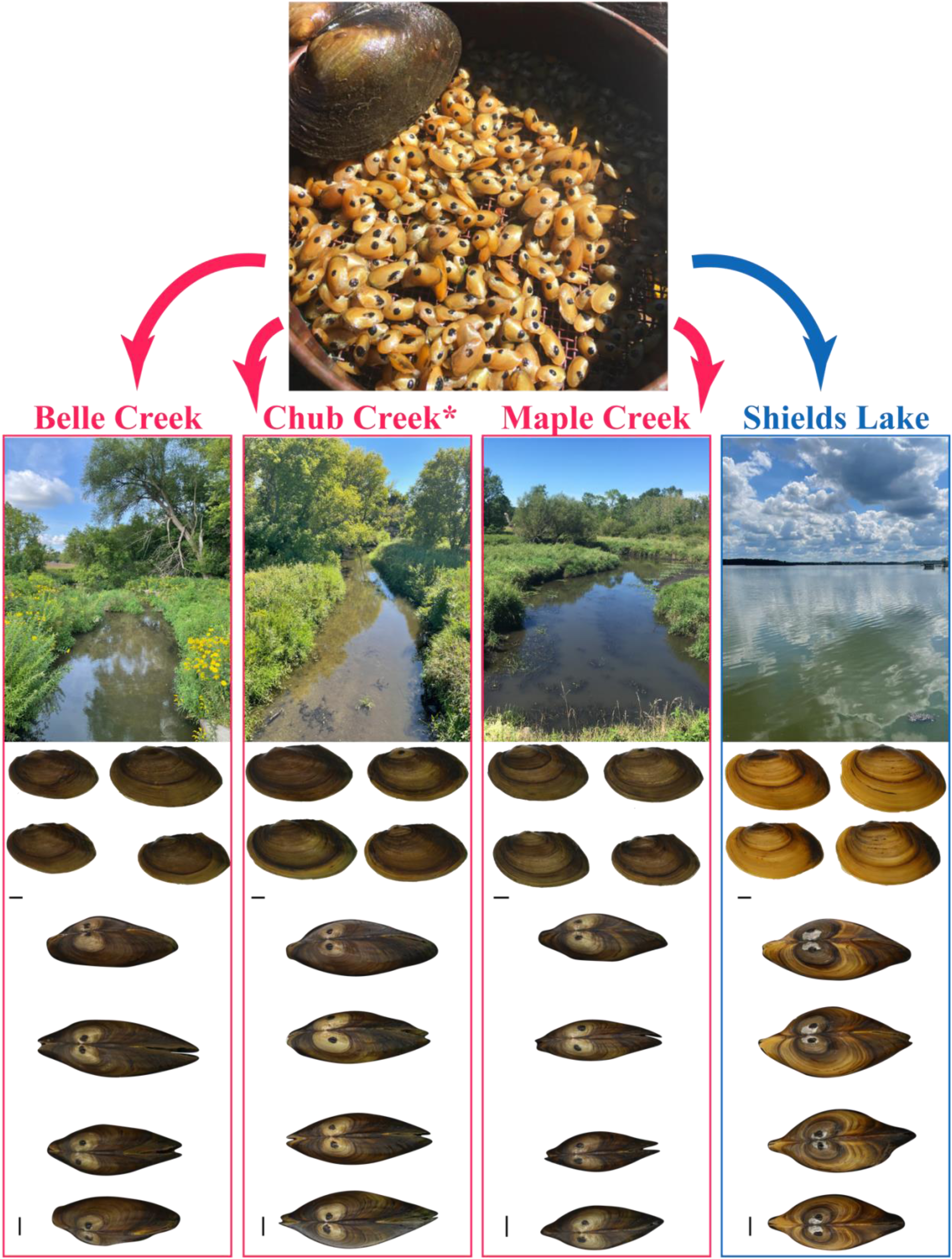
Layout of experimental design and shell morphology of recaptured siblings for four field sites including two habitats. Top photo shows marked siblings at three months of age immediately prior to field release with an adult *P. grandis* for scale. Second row shows each field site at time of recapture. Third and fourth rows show four representative mussels from each recapture site: lateral view (top) and dorsal view (bottom). Black bars equal 1 cm for each image.

### 5.2 Genetic variation and paternity testing

Single nucleotide polymorphisms (SNPs) were obtained for 39 samples (1 mother, 38 recaptured offspring). Genetic distances, F_ST_, between sites were low, ranging from 0.002-0.02 (Supplemental Table 2) and effectively zero between habitats (–0.00001; N=19 in each habitat, mother omitted from analysis). COLONY identified 27 (unsampled) fathers for the 38 recaptured mussels. Eight clusters of full siblings were identified containing 2-3 samples each. Five clusters contained individuals collected from both stream and lake habitats and three clusters contained individuals exclusively from Shields Lake (Figure 5).

### 5.3 Statistical analysis

#### 5.3.1 Q1: Do mussels reared at different sites have different shell morphology?

We found rearing site had a significant effect (Table 1; Figure 4a-b) on shell height and width while accounting for covariation with shell length showing that some aspect of the environment induces morphological change. We found significant differences in shell length indicating that shell growth differed among sites. Additionally, the interaction between site and shell length was significant for the shell width model excluding the zoo population and for both shell height and width for models including the zoo population. Significant interaction effects indicate site differences in the slopes of shell width∼length and shell height∼length, meaning allometric differences between rearing sites. Although we did not quantify shell coloration, we observed different shell coloration across the four sites (Figure 3).

**Figure 4:**
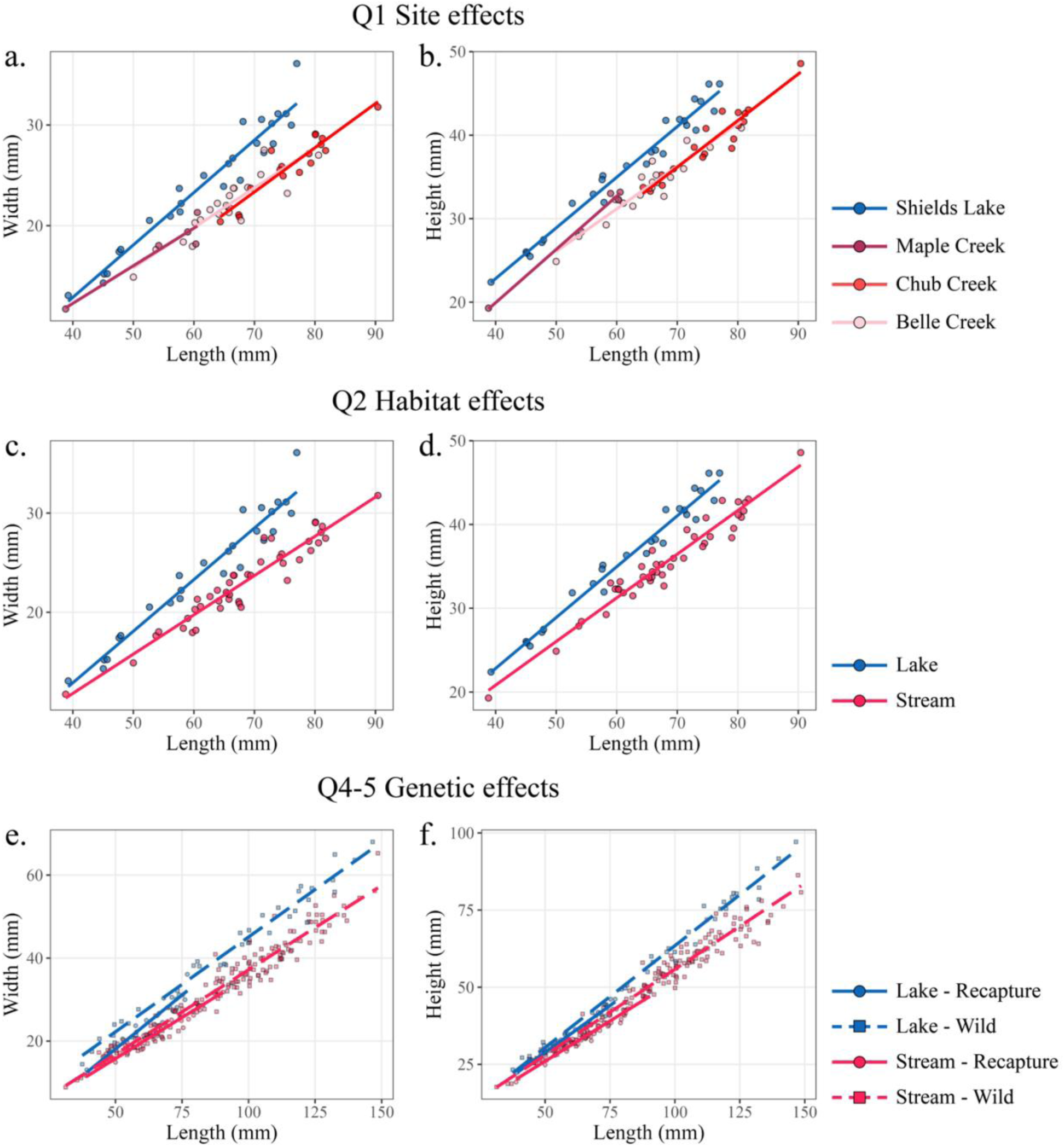
Linear regressions of shell width (left panels) and shell height (right panels) as a function of shell length (i.e. growth trajectories). Top row (a. & b.): recaptured mussels reared at each site have a separate regression line. Second row (c. & d.): growth regressions for recaptured mussels reared in each habitat type. Bottom row (e. & f.): growth regressions for wild and recaptured populations in each habitat type.

**Figure 5:**
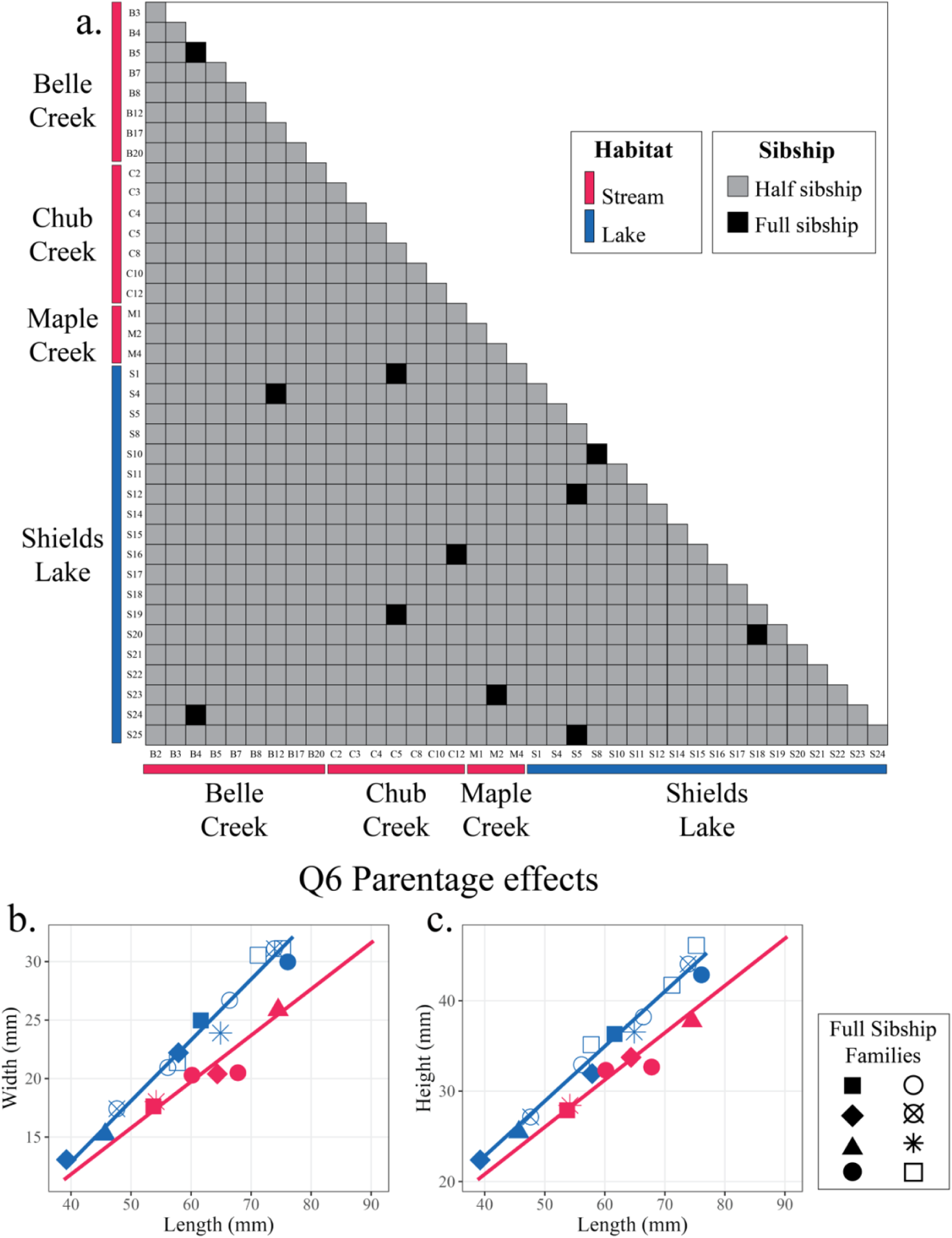
Sibship assignments from COLONY by habitat type and site for 38 sequenced individuals (a.). Eight full sibship families were identified and their shell shapes including shell width (b.) and shell height (c.) are plotted. Regression lines (b. & c.) are for the full 70 recaptured individuals (identical to Figure 4c-d). All full sibship families had probabilities >0.99. In total, 27 fathers were identified in the COLONY analysis but only individuals with at least one full sibling are plotted in b-c.

**Table 1:**
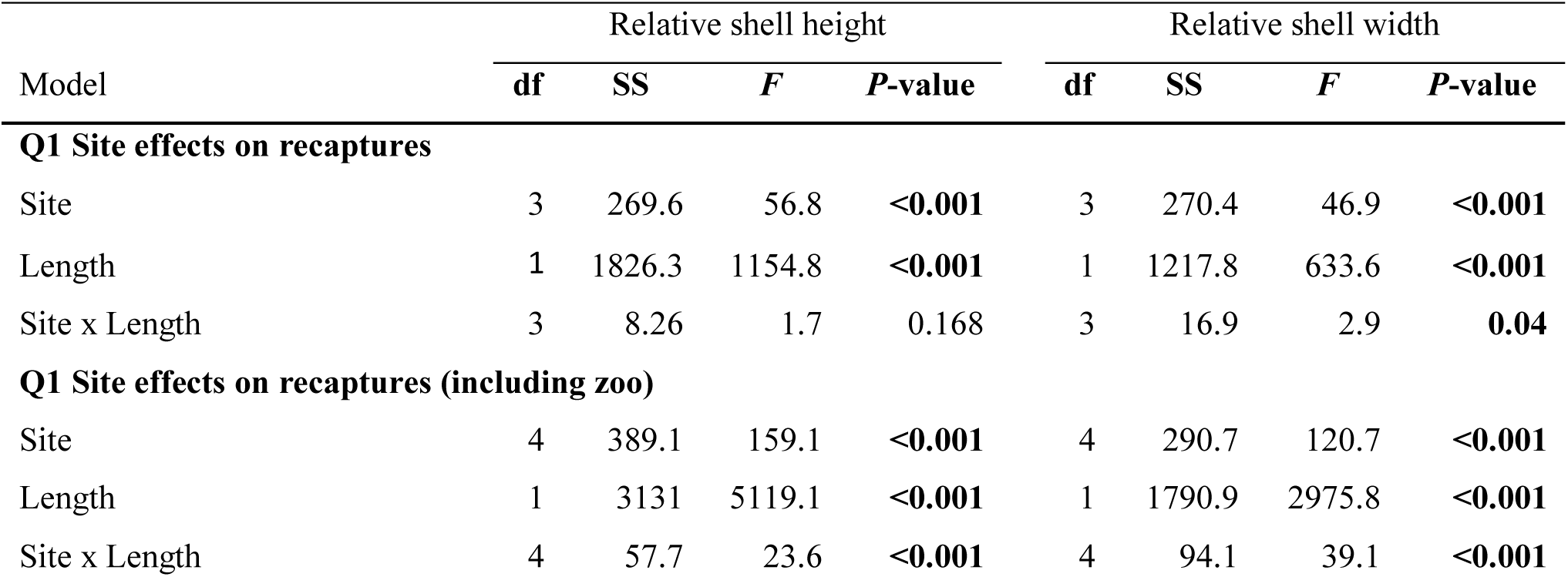
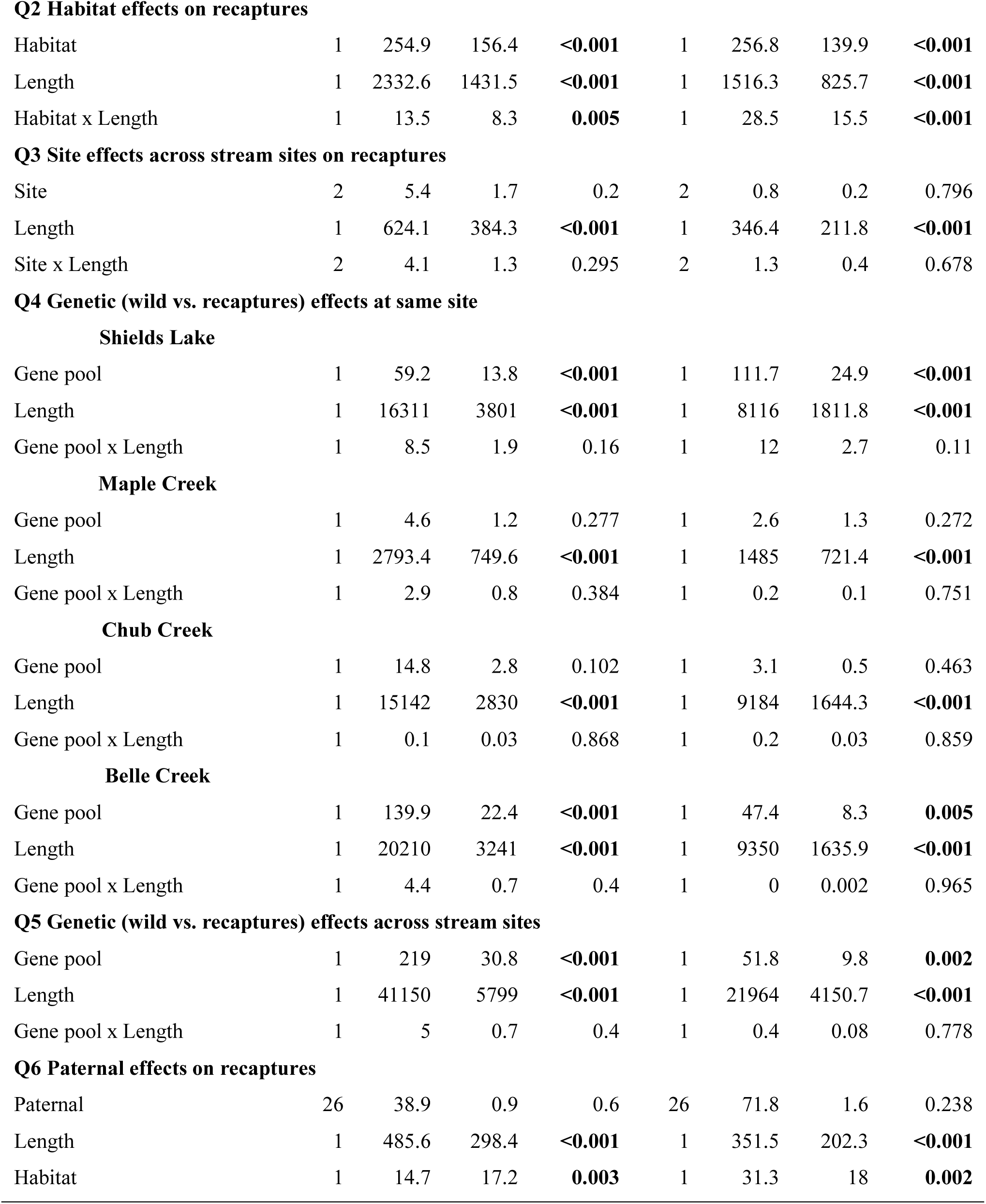
Results of ANCOVAs (questions 1-6) including degrees of freedom (df), sum of squares (SS), F statistic (*F*), and p-value. P-values ≤ 0.05 are bolded.

| Model | Relative shell height |  |  |  | Relative shell width |  |  |  |
| --- | --- | --- | --- | --- | --- | --- | --- | --- |
|  | df | SS | *F* | *P*-value | df | SS | *F* | *P*-value |
| **Q1 Site effects on recaptures** |  |  |  |  |  |  |  |  |
| Site | 3 | 269.6 | 56.8 | **<0.001** | 3 | 270.4 | 46.9 | **<0.001** |
| Length | 1 | 1826.3 | 1154.8 | **<0.001** | 1 | 1217.8 | 633.6 | **<0.001** |
| Site x Length | 3 | 8.26 | 1.7 | 0.168 | 3 | 16.9 | 2.9 | **0.04** |
| **Q1 Site effects on recaptures (including zoo)** |  |  |  |  |  |  |  |  |
| Site | 4 | 389.1 | 159.1 | **<0.001** | 4 | 290.7 | 120.7 | **<0.001** |
| Length | 1 | 3131 | 5119.1 | **<0.001** | 1 | 1790.9 | 2975.8 | **<0.001** |
| Site x Length | 4 | 57.7 | 23.6 | **<0.001** | 4 | 94.1 | 39.1 | **<0.001** |
| **Q2 Habitat effects on recaptures** |  |  |  |  |  |  |  |  |
| Habitat | 1 | 254.9 | 156.4 | **<0.001** | 1 | 256.8 | 139.9 | **<0.001** |
| Length | 1 | 2332.6 | 1431.5 | **<0.001** | 1 | 1516.3 | 825.7 | **<0.001** |
| Habitat x Length | 1 | 13.5 | 8.3 | **0.005** | 1 | 28.5 | 15.5 | **<0.001** |
| **Q3 Site effects across stream sites on recaptures** |  |  |  |  |  |  |  |  |
| Site | 2 | 5.4 | 1.7 | 0.2 | 2 | 0.8 | 0.2 | 0.796 |
| Length | 1 | 624.1 | 384.3 | **<0.001** | 1 | 346.4 | 211.8 | **<0.001** |
| Site x Length | 2 | 4.1 | 1.3 | 0.295 | 2 | 1.3 | 0.4 | 0.678 |
| **Q4 Genetic (wild vs. recaptures) effects at same site****Shields Lake** |  |  |  |  |  |  |  |  |
| Gene pool | 1 | 59.2 | 13.8 | **<0.001** | 1 | 111.7 | 24.9 | **<0.001** |
| Length | 1 | 16311 | 3801 | **<0.001** | 1 | 8116 | 1811.8 | **<0.001** |
| Gene pool x Length | 1 | 8.5 | 1.9 | 0.16 | 1 | 12 | 2.7 | 0.11 |
| **Maple Creek** |  |  |  |  |  |  |  |  |
| Gene pool | 1 | 4.6 | 1.2 | 0.277 | 1 | 2.6 | 1.3 | 0.272 |
| Length | 1 | 2793.4 | 749.6 | **<0.001** | 1 | 1485 | 721.4 | **<0.001** |
| Gene pool x Length | 1 | 2.9 | 0.8 | 0.384 | 1 | 0.2 | 0.1 | 0.751 |
| **Chub Creek** |  |  |  |  |  |  |  |  |
| Gene pool | 1 | 14.8 | 2.8 | 0.102 | 1 | 3.1 | 0.5 | 0.463 |
| Length | 1 | 15142 | 2830 | **<0.001** | 1 | 9184 | 1644.3 | **<0.001** |
| Gene pool x Length | 1 | 0.1 | 0.03 | 0.868 | 1 | 0.2 | 0.03 | 0.859 |
| **Belle Creek** |  |  |  |  |  |  |  |  |
| Gene pool | 1 | 139.9 | 22.4 | **<0.001** | 1 | 47.4 | 8.3 | **0.005** |
| Length | 1 | 20210 | 3241 | **<0.001** | 1 | 9350 | 1635.9 | **<0.001** |
| Gene pool x Length | 1 | 4.4 | 0.7 | 0.4 | 1 | 0 | 0.002 | 0.965 |
| **Q5 Genetic (wild vs. recaptures) effects across stream sites** |  |  |  |  |  |  |  |  |
| Gene pool | 1 | 219 | 30.8 | **<0.001** | 1 | 51.8 | 9.8 | **0.002** |
| Length | 1 | 41150 | 5799 | **<0.001** | 1 | 21964 | 4150.7 | **<0.001** |
| Gene pool x Length | 1 | 5 | 0.7 | 0.4 | 1 | 0.4 | 0.08 | 0.778 |
| **Q6 Paternal effects on recaptures** |  |  |  |  |  |  |  |  |
| Paternal | 26 | 38.9 | 0.9 | 0.6 | 26 | 71.8 | 1.6 | 0.238 |
| Length | 1 | 485.6 | 298.4 | **<0.001** | 1 | 351.5 | 202.3 | **<0.001** |
| Habitat | 1 | 14.7 | 17.2 | **0.003** | 1 | 31.3 | 18 | **0.002** |

#### 5.3.2 Q2: Do mussels reared at different habitats have different shell morphology?

When aggregating recaptured individuals into stream or lake habitat types, we found a significant difference in both shell height and width between habitats (Table 1; Figure 4c-d). Significant differences in shell length and interaction between habitat and shell length were found for both shell height and width models. Qualitatively, shell color was different between lake and stream rearing sites with lake individuals colored bright yellow and stream individuals a dark chestnut (Figure 3).

#### 5.3.3 Q3: Do mussels reared at different sites but the same habitat type have different shell morphology?

Our ANCOVA models failed to find shell morphology differences between the three stream sites (Table 1). This result was supported by the pairwise *post hoc* tests which found all pairwise combinations of stream sites produced no significant differences in shell morphology (Supplemental Table 3). However, shell length did differ between stream sites.

#### 5.3.4 Q4: Do recaptured mussels have different shell morphology than wild mussels reared at the same site?

Morphological differences between recaptured and wild reared mussels were site specific: no shell shape differences were found at Maple and Chub Creek but relative shell height and width were significantly different at both Shields Lake and Belle Creek (Table 1; Figure 4e-f). Shell length was significantly different at each site as recaptured mussels were being compared against wild mussels of all age classes.

#### 5.3.5 Q5: Do recaptured mussels have different shell morphology than wild mussels reared at the same habitat type?

Although this test was only performed for the stream habitat type, the results followed those of question 4. When the ANCOVA included Belle Creek mussels it found significant morphological differences between recaptured and wild mussels in stream habitats (Table 1; Figure 4e-f).

#### 5.3.6 Q6: Does paternity affect shell morphology?

While accounting for covariation with shell length and habitat, we found no paternal effect on either shell width or height (Table 1; Figure 5b-c). There was also significant effects of habitat type on both shell width and height, a likely consequence of three of the eight full sibship families containing individuals exclusively from Shields Lake. Parentage did have a significant effect on shell length.

## 6. Discussion

The results of our field experiment show extreme shape and color divergence among recaptured mussels reared in different sites and habitat types (Table 1; Figure 3-4) but no genetic difference between recaptured mussels from stream and lake habitats (F_ST_=-0.00001) and no parentage effect on shell morphology (Table 1, Q6), thereby revealing phenotypic plasticity as the primary mechanism producing ecophenotypic variation in *P. grandis*. The combined evidence of shell shape differences between sites (Q1) and habitat types (Q2) as well as a lack of differentiation across the three stream sites (Q3) shows that plasticity in shell shape responded most robustly and predictably to habitat type: lake versus stream environments (Figure 4b-c). Ecophenotypic expressions of recaptured *P. grandis* match previous field observations (Ortmann 1919; Clarke 1973) and in some cases matched the morphology of wild captured mussels reared at the same site (Table 1; Figure 4e-f). Across the three stream sites, shells were elongated in the anterior-posterior orientation, laterally deflated, and periostracum (i.e. outermost shell layer) color was dark brown. Conversely, shells at Shields Lake were circular in both the longitudinal (anterior-posterior) and axial (width) planes with a bright yellow periostracum (Figure 3). Mean shell height and width of recaptured mussels mirrored that of wild collected mussels at Maple and Chub Creek but morphology between wild and recaptured mussels was distinct at both Belle Creek and Shields Lake (Table 1, Q4). This mismatch between experimental and wild mussels suggests that genetic differentiation may still play a role in ecophenotypic outcomes. These sites were the only sites where high sample sizes of first– and second-year-old wild mussels were collected and because recaptured mussels spent most of their first growing season (May-August 2020) in laboratory conditions. Therefore, this mismatch may reflect a developmental lag to their new environment (see regression slopes in Figure 4e-f) rather than genetic differentiation. Lastly, of 38 individuals sequenced from the same maternal brood, 27 fathers were identified (Figure 5a). This high degree of multiple paternity, where many males contribute genetically to a single reproductive event, may be an important mechanism to harbor high genetic variation within mussel populations.

### 6.1 Environmentally mediated phenotypes

Phenotypic plasticity is expected to evolve in lineages that experience high spatial and temporal variation in environmental conditions and have high dispersal capabilities (Hendry 2016; Pfennig 2021). Temporary parasites, who rely on their hosts for dispersal may uniquely fit these criteria. Freshwater mussels, with few exceptions, are obligate parasites during the larval life-stage, relying on an aquatic vertebrate (typically fish) host for dispersal. The mobility of the host is the primary determinant of dispersal distance and habitat use generation-to-generation (Terui et al. 2017; Watters 1992). Host specificity varies greatly across freshwater mussels (Hewitt et al. 2019) including host specialists (Sietman et al. 2017; 2018) and host generalists including *P. grandis* who can successfully transform on over 30 fish species from 11 taxonomic families (Hopper et al. 2023). Intuitively, host generalism would increase environmental uncertainty and dispersal variance. *Pyganodon grandis* uses large, highly migratory fishes including *Micropterus nigricans* (Largemouth bass) and *Alosa chrysochloris* (Skipjack herring) but also smaller, more localized fishes including *Fundulus diaphanus* (Banded killifish) and *Etheostoma nigrum* (Johnny darter) (Lefevre and Curtis 1910; Surber 1913; Trdan and Hoeh 1982). Therefore, host generalism may be a predisposition to phenotypic plasticity as has been proposed in other systems (Brown et al. 2012; Leggett et al. 2013). Alternatively, host generalism may be a product of plasticity as adaptive phenotypic plasticity would increase the potential habitats occupied by a species and thus increase interactions with potential hosts in novel habitats. With their high variance in host specificity and documented phenotypic plasticity here, freshwater mussels may be a uniquely suited system to test these causal relationships.

Habitat type was the biggest determinant of shape variation for both recaptured and wild caught *P. grandis* (Figure 4). However, the specific environmental cue(s) initiating phenotypic expression is unclear. Hydrology is the most obvious environmental difference between lake and stream field sites and quantification of within-species ecophenotypic relationships between morphology and water velocity (Balla and Walker 1991), flow rate (i.e. discharge) (Graf 1998; Keogh et al. In Press), Reynolds number (i.e. hydraulic energy) (Simeone et al. 2022), and Strahler stream order (Ortmann 1920; Keogh et al. In Press) suggest that freshwater mussels reliably respond to hydrologic changes. However, the zoo population, reared in basket enclosures within a lake, had drastically different morphology with more ‘stream-like’ relative heights and widths (Supplementary Figure 1). These mussels were reared alongside baskets of other freshwater mussel species and to ensure adequate dissolved oxygen, air pumps were used surrounding the baskets. Further, the baskets were densely packed with upwards of ∼700 mussels with an average length of 42 mm in a ∼1.5 meters^2^ area. Hydrologic agitations caused by either neighboring mussels or air pumps could have been enough to falsely trigger ‘stream-form’ development. Importantly, there was very little morphological variance among the 305 individuals measured at the zoo showing that they were all responding to identical signals. In addition to hydrology, substrate type (Hinch et al. 1986) and turbidity (Tuttle-Raycraft and Ackerman 2020) have both been shown to elicit plastic phenotypic responses in freshwater mussels. Yet, anecdotally, substrate was not noticeably different between Shields Lake and the stream sites. At stream sites, mussels were collected from clay, silt, sand, and gravel and at Shields Lake, silt and sand. Turbidity and other water quality parameters are unlikely determinants of shell shape as ‘lake-form’ and ‘stream-form’ phenotypes can be collected meters away within the same contiguous waterbody (e.g. above and below dams, Haag 2012). More common garden experiments, preferably in laboratory conditions where environmental variation can be more finely manipulated, are necessary to illuminate the specific environmental cue(s) initiating shape change.

Phenotypic plasticity can produce neutral, maladaptive, or adaptive morphologies. Whether fitness advantages are tied to shell shape variation in *P. grandis* remains an unanswered question but the shared phenotypic expression of shell inflation in lake habitats and shell deflation in stream habitats across multiple divergent lineages (Grier 1920) suggests phenotypic plasticity is adaptive in *P. grandis*. Watters (1994) hypothesized that shell inflation in lakes is an adaptation for buoyancy in fine substrates (e.g. silt). Shell inflation increases surface area perpendicular to the burrowing axis and life position, which along with thin shells, may decrease sinking in low-density substrates. As noted above, fine substrates can occur in stream habitats as well but Watters (1994) suggested shell inflation is deleterious in stream habitats because streamflow can dislodge inflated shells. Alternatively, Eager (1978) proposed that the ‘stream-form’ and ‘lake-form’ phenotypic gradient reflects a tradeoff between stability in hydrodynamically demanding stream environments versus metabolic efficiency. Released from the physical constraints of streamflow, shell growth and morphology in lake populations may reflect an optimization of energy allocation towards internal soft anatomy involved in feeding, respiration, and reproduction rather than shell secretion. An optimal way to reduce surface area and thus shell production while increasing internal volume is to become more spherical (Eager 1979), as the ‘lake-form’ clearly does (Figure 3). Lastly, the spherical shape divergence in lake habitats may be an anti-predator adaptation to more quickly escape the gape of molluscivorous fishes. Despite releasing more animals in lakes and having more lake sites than streams, we only recaptured mussels from a single lake site. Our recapture efforts may reflect lower survival in lake versus stream habitats with predation being the underlying cause. Additional physiological and functional experiments are needed to illuminate the adaptive significance of shell forms.

### 6.2 Conservation implications

In the face of widespread imperilment (Haag 2019; Haag and Williams 2014; Hornbach et al. 2018; Ollard and Aldridge 2023), the identification of phenotypic plasticity in freshwater mussels has significant conservation implications. First, extreme ecophenotypy in species, like *P. grandis*, reflects their evolutionary potential. Regardless if this variation is adaptive, it illustrates that plastic freshwater mussel species have the ability to respond to changing environments: a trait that may provide resilience to anthropogenically modified habitats and climate change (Reed et al. 2011). Second, the identification of phenotypic plasticity rather than genetic differentiation as the mechanism for ecophenotypic patterns is highly convenient for conservation managers. The most popular tool used to combat mussel imperilment is captive-rearing propagation (FMCS 2016; Haag and Williams 2014). Propagation depends on field collection of gravid individuals, captive-rearing and culture of juveniles, and release of reared individuals, frequently in different localities than the collected parents (Gum et al. 2011). The identification of phenotypic plasticity in a given lineage allows managers some flexibility in release location and ontogenetic timing as plastic species like *P. grandis* are highly acclimatized to environmental heterogeneity generation-to-generation and possibly within lifetimes. Cultured mussels spend variable amount of time in captivity and the timing of release into novel, wild localities may be particularly important if developmental plasticity is fixed beyond a certain life point (M. Pletta pers. comm.). However, no discernable variation in shape was detected when mussels were initially released into wild sites when three months old, but the shape divergence observed after two years shows that developmental plasticity does not become fixed in *P. grandis*, at least not early in life. In addition to plasticity, multiple paternity and thus high genetic variation in a single brood (Figure 5a) suggests propagated and released individuals are not homogenizing genetic variation of natural populations (Inoue et al. 2023). The prevalence of propagation programs in North America are now being leveraged for basic science, including the research conducted here but also in ecotoxicology (Buczek et al. 2017, Popp et al. 2018), thermal tolerance (Pandolfo et al. 2010; Archambault et al. 2013), and life-history (Lefevre and Curtis 1910; Coker et al. 1921; Howard 1922; Sietman et al. 2017; 2018), which may be equally valuable to mussel conservation as stocking.

### 6.3 Conclusion

Experimental results from this study and others (Hinch et al. 1986; Tuttle-Raycraft and Ackerman 2020) show that freshwater mussels can conditionally express their anatomical features in response to environmental cues and phenotypic plasticity is likely a ubiquitous feature of mussels given strong ecophenotypic variation in other species (Grier 1920; Ortmann 1920; Ball 1922; Agrell 1948; Graf 1998; Balla and Walker 1991; Zieritz and Aldridge 2009; Keogh et al. In Press). Given its high and predictable morphological variation, *Pyganodon grandis* may be an ideal model system to study phenotypic plasticity and moreover, the functional genomics of shell shape in bivalves. We reared nearly ∼7,000 individuals to three months of age from a single mother and found they grew quickly and were robust to environmental changes. Additional investigation is needed to identify the environmental cues and adaptive significance of shell shape variation. Integrative and experimental inferences such as combining transcriptomics, morphometrics, and functional experiments are warranted to further understand phenotypic plasticity in freshwater mussels.

## 7. Acknowledgments

We thank Megan Bradley and Beth Glidewell for discussions on the propagation of *Pyganodon grandis*, Camille Will, Owen Bachhuber, and Zoe Sax for assistance in the field and mussel husbandry, and Keith Barker for methodological suggestions. The Minnesota Supercomputing Institute (MSI) at the University of Minnesota was used to assemble genomic data (http://www.msi.umn.edu).

## 10. Data Availability

Morphological data, R scripts for analyses, and COLONY input and results can be found at https://github.com/seanmkeogh/Pgrandis_plasticity/tree/main. Demultiplexed Illumina sequence data will be uploaded to NCBI SRA upon acceptance.

## 11. Authors contributions

Conceptualization: S.M.K., B.J.M., B.E.S., A.M.S.; Methodology: S.M.K.; Software: S.M.K.; Validation: S.M.K.; Formal analysis: S.M.K.; Investigation: S.M.K., B.J.M., L.M.O., M.E.P., A.E.S., Z.K.S., Z.A.S., A.J.F., B.E.S.; Resources: S.M.K., B.J.M., B.E.S., A.M.S.; Data curation: S.M.K.; Writing – original draft preparation: S.M.K.; Writing – review and editing: S.M.K., B.J.M., L.M.O., A.J.F., B.E.S., A.M.S.; Visualization: S.M.K.; Supervision: S.M.K., B.J.M., B.E.S., A.M.S.; Project administration: S.M.K., B.J.M., B.E.S.; Funding acquisition: S.M.K., B.J.M., A.M.S.

